# Movement of transposable elements contributes to cichlid diversity

**DOI:** 10.1101/2020.02.26.961987

**Authors:** Karen L. Carleton, Matt Conte, Milan Malinsky, Sri Pratima Nandamuri, Ben Sandkam, Joana I Meier, Salome Mwaiko, Ole Seehausen, Thomas D Kocher

**Affiliations:** Department of Biology, University of Maryland, College Park MD 20742 USA; Wellcome Sanger Institute, Cambridge, UK; Zoological Institute, University of Basel, Basel, Switzerland; Aquatic Ecology and Evolution, Institute of Ecology and Evolution, University of Bern, 3012 Bern, Switzerland; Department of Fish Ecology and Evolution, Centre for Ecology, Evolution & Biogeochemistry, Eawag: Swiss Federal Institute of Aquatic Science and Technology, 6047 Kastanienbaum, Switzerland; Computational and Molecular Population Genetics Lab, Institute of Ecology and Evolution, University of Bern, 3012 Bern, Switzerland

**Author notes:** John A Moran Center, University of Utah, Salt Lake City UT USA. Department of Zoology, University of British Columbia, Vancouver, BC, Canada. Department of Biology, University of Cambridge, Cambridge UK.

## Abstract

African cichlid fishes are a prime model for studying the mechanisms of speciation. Despite the development of extensive genomic resources, it has been difficult to determine which sources of genetic variation are responsible for variation in cichlid phenotypes. Cichlids have some of the largest known shifts in vertebrate visual sensitivity. These shifts arise mainly from the differential expression of seven cone opsin genes. By mapping expression quantitative trait loci (eQTL) in intergeneric crosses of Lake Malawi (LM) cichlids, we have thus far identified four causative genetic variants that correspond to indels in the promoters of either key transcription factors or of the opsin gene itself. Here we show that these indels are caused by the movement of transposable elements (TEs). These precise indels are not found outside of LM, suggesting that these TEs are recently active and are segregating within the Malawi cichlid lineage. A similar indel has arisen independently outside of LM at one locus, suggesting that some locations are primed for TE insertion and the resulting indels. Increased TE mobility may be associated with interspecific hybridization, which disrupt mechanisms of TE suppression. Overall, our study suggests that TEs may contribute to key regulatory changes, and may facilitate rapid phenotypic change and possibly speciation in African cichlids.

## Introduction

The genomic era promised to unlock the molecular links between genotype and phenotype, including the evolution of new species. Questions of evolutionary predictability have focused on whether new phenotypes result from changes in coding sequence or gene regulation (Hoekstra and Coyne 2007, Carroll 2008, Stern and Orgogozo 2008). In addition, there has been a strong focus on single nucleotide polymorphisms (SNPs) during the search for highly selected regions, magic genes, and islands of speciation (Turner and Hahn 2010, Servedio et al. 2011, Malinsky et al. 2015, Malinsky et al. 2018, Svardal et al. 2019). However, evolution likely occurs through more than changes to single DNA bases. Structural rearrangements, such as insertions and deletions (indels; Mills et al. 2006, Green et al. 2010), inversions, gene duplications (Ohno 1970, Cortesi et al. 2015), or whole genome duplications (Otto and Whitton 2000, Crow et al. 2006) can be critical to generating evolutionary novelty.

The approximately 1500 species of African cichlid fishes are a textbook example of adaptive radiations, characterized by phenotypic change in a rapidly speciating lineage (Kocher 2004, Malinsky and Salzburger 2016). Cichlids differ in many extensively studied and ecologically important phenotypes, including examples such as jaw morphology (Albertson et al. 2003), color patterns (Seehausen 1996, Danley and Kocher 2001, Allender et al. 2003, Konings 2007), sex determination (Roberts et al. 2009, Gammerdinger and Kocher 2018), and parental care (Barlow 2000, Sefc 2011).

To understand the genetic basis of this diversity, the genomes of five species were sequenced by the Cichlid Genome Consortium (Brawand et al. 2014). This included one species from each of the three Great African Lakes (Malawi, Tanganyika and Victoria), as well as a recent and a more ancestral riverine species. Based on the genomic analyses, at least five mechanisms were suggested to explain cichlid diversity. First was an acceleration of sequence evolution, including higher dN/dS ratios, in genes controlling development, pigmentation and vision. Second was a 4.5-6x increase in the rate of gene duplications in the ancestor of the rapidly radiating lacustrine cichlid lineages, when compared to non-cichlid fish such as stickleback and zebrafish. Third, there were 625 regulatory regions, which showed accelerated evolution within the East African lake species. Fourth, there were 40 gains and 9 losses of miRNAs. Finally, there was evidence for transposable element insertions that were associated with changes in gene expression. Although all of these mechanisms are likely to contribute to cichlid diversity, none of them have yet been tied to specific phenotypes. The genetic architecture of some ecologically important cichlid traits, such as body shape, is likely to be highly polygenic. For such traits, the contributions of numerous loci of small effect are very difficult to determine. However, the link between genotypes and phenotype is easier to make for traits with simpler genetic architectures dominated by large effect (sometimes even mendelian) loci, such as pigmentation patterns (Santos et al. 2014, Kratochwil et al. 2018), some aspects of jaw development (e.g. Roberts et al. 2011, Parsons et al. 2014, Conith et al. 2018), or various adaptations of the visual system (e.g. Sugawara et al. 2005, Malinsky et al. 2015, Carleton et al. 2016, Malinsky et al. 2018).

In this study, we focus on the evolution of cichlid visual systems. Visual systems have the advantage that opsin genes are already known to be the key genes shaping visual sensitivity phenotypes. Opsin proteins combine with a chromophore, such as 11-cis retinal, to produce visual pigments that absorb light (Yokoyama 2008). The sequence of the opsin protein is key to tuning visual pigment sensitivity. Because of the importance of visual systems to survival, they are likely under strong natural selection (Davies et al. 2012, Goldsmith 2013).

Microspectrophotometric measurements of cichlid rod and cone cells suggest that cichlid visual systems are highly variable among species (Levine et al. 1979, van der Meer and Bowmaker 1995, Jordan et al. 2006). Cichlids have short wavelength single cones and longer wavelength double cones. Species can differ in whether their single cones are ultraviolet, violet or blue sensitive with peak sensitivities varying by up to 90 nm (Jordan et al. 2006, Carleton 2009). Double cones also vary by 30 to 50 nm, with closely related species showing remarkable shifts in peak sensitivity.

In previous work, we have started to identify the genetic mechanisms shaping visual diversity. Cichlids have seven cone opsin genes, which belong to the four vertebrate classes: very short wavelength sensitive 1 (SWS1), short wavelength sensitive 2 (SWS2A and SWS2B), rhodopsin like (RH2Aa, RH2Ab and RH2B) and long wavelength sensitive (LWS). Protein expression confirmed that these seven genes produce distinct visual pigments with peak sensitivities distributed across the spectrum from ultraviolet to red wavelengths (Parry et al. 2005, Spady et al. 2006). Comparisons across different species suggest that opsin sequences may evolve rapidly and adaptively, but sequence differences contribute relatively small spectral shifts (Spady et al. 2005, Sugawara et al. 2005, Seehausen et al. 2008, Nagai et al. 2011, Malinsky et al. 2018). Therefore, while opsin protein sequences are important, they are not the primary driver of the larger shifts in visual sensitivity (Carleton and Kocher 2001, Hofmann et al. 2009).

Differential expression of opsin genes is key to cichlid visual diversity. Adults are typically trichromatic, expressing three cone opsins more than others (Carleton 2009, Carleton et al. 2016). The three common gene combinations are the short (*SWS1, RH2B, RH2A*), medium (*SWS2B, RH2B, RH2A*) and long (*SWS2A, RH2A, LWS*) visual palettes (with *RH2A* signifying either of the highly similar *RH2Aα* and *RH2Aβ* genes). Although many species only express one palette throughout life, some species progress from the short to medium to long combinations through development (Carleton et al. 2008, O’Quin et al. 2011).

To identify the loci underlying these differences in opsin expression, we made genetic crosses between species expressing different palettes. The causative genetic factors we found are typically not in cis. Instead we found several expression quantitative trait loci (eQTL) acting in trans to the opsin genes (O’Quin et al. 2012, Nandamuri et al. 2018). Using fine mapping in crosses, and association mapping in natural populations, we have identified the causative genes as well as putative mutations that might underlie these changes. Retinal homeobox 1 (*Rx1*; Schulte et al. 2014) and microphthalmia associated transcription factor a (*Mitfa*; Nandamuri 2018) are trans factors associated with changes in the expression of the *SWS2A* opsin gene. A 413bp deletion that is 2.5 kb upstream of the Rx1 translational start site causes a decrease in *SWS2A* expression in Lake Malawi cichlids. It explains 62% of the variance in *SWS2A* expression across over 50 species (Schulte et al. 2014). An insertion in intron 1 of *Mitfa* is correlated with an increase in *SWS2A* expression, though it has a smaller effect than *Rx1* (Nandamuri 2018). T-box 2a (*Tbx2a*) is a trans factor associated with expression of the *LWS* opsin (Sandkam et al. in press). We have shown that *Tbx2a* binds to regulatory regions for both LWS and RH2 and acts to switch between these opsins. A 967bp deletion that is 13.5kb upstream of its translational start site causes the shift from LWS to RH2 expression in one species in our cross. In addition to these three regulatory mutations near transcription factors, there is one cis regulatory change associated with changes in the expression of *SWS1* (Nandamuri et al. 2018). This deletion removes several conserved regulatory elements shutting off *SWS1* expression. Therefore, in each of these cases, we have found a regulatory indel that either removes (*Rx1, Tbx2a, SWS1*) or adds (*Mitfa*) a critical regulatory region. These indels alter either the expression of the critical transcription factor affecting opsin expression, or directly alters the promoter sequence of the opsin itself (*SWS1*).

In the current study, we characterize the evolutionary origins and mechanisms of these four mutations. We find that they involve either insertions or deletions of significant size (400-2000 bp). The boundaries of these indels largely correspond to transposable elements. By examining species within and outside the Malawi flock including from the sister Lake Victoria lineage, we discovered these indels are recent and, with one exception, specific to the cichlids of Lake Malawi. Further, they seem linked to an increase in the number of copies of particular TE families. We suggest that the movement of transposable elements generates sizeable indels that modify important regulatory regions. TE movement may be an unappreciated but key mechanism underlying cichlid diversity.

## METHODS

### Indel analysis

We focus on the four mutations identified in our previous QTL studies. These include the regulatory regions for three transcription factors, *Rx1, Tbx2a, Mitfa* and the *SWS1* opsin. Potentially causative mutations in these regulatory regions were first characterized in genetic crosses between Lake Malawi cichlids. The loci for *Rx1, Tbx2a* and *Mitfa* were identified in a cross between *Tramitichromis intermedius* and *Aulonocara baenschi*. The *SWS1* locus was characterized in a cross between *Metriaclima* ‘mbenji’ and *A baenschi*. These loci were then compared to the genome of *Metriaclima zebra* as well as genomes of outgroups to Lake Malawi including *Astatotilapia burtoni*, *Pundamilia nyererei* and *Oreochromis niloticus* sequenced as part of the cichlid genome project (Brawand et al. 2014). This determined whether a locus was an insertion or a deletion relative to the outgroup species, as well as its relative size. Inserted sequences were analyzed using Repbase (Kohany et al. 2006) using the CENSOR website (https://girinst.org/censor/index.php) to determine if they corresponded to known transposable elements.

### Origin and phylogenetic diversity of indel sequences

To identify the phylogenetic origin and to estimate the age of these indels, we searched for them across species within and outside of Lake Malawi using a combination of PCR and whole genome sequences. In total, 209 species and 235 individuals were surveyed. This included examining these regions in the five cichlid genome project species (Brawand et al. 2014). Next, single individuals of 53 Lake Malawi species where we have quantified opsin gene expression (Hofmann et al. 2009) were screened by PCR (Supp Table 1). In addition, we used PCR to screen for the *Rx1* and *Mitf* indels in three species (n=1) from Lake Malawi, one from Lake Chilingali (a lake very close to Lake Malawi) and five populations (n=1-2) of *Astatotilapia calliptera* from nearby rivers, which group within the Lake Malawi rock dwelling clades (Joyce et al. 2011; individuals and species are listed in Supp Table S2). We also searched newly sequenced genomes of 103 Lake Victoria species (86 from Lake Victoria proper, 2 from Lake Nabugabo and 15 from Lake Kyoga) as well as 11 riverine outgroup species. Lake Victoria species were Illumina sequenced and the reads mapped onto the *Pundamilia nyererei* genome. Variants were called with GATK to make VCF tracks for viewing in the Integrative Genome Viewer (Robinson et al. 2011, Thorvaldsdottir et al. 2013). Finally, we searched 52 additional genomes including 15 species from Lake Malawi and 37 outgroups. These taxa were sequenced to ~15x coverage on the Illumina platform and the raw reads were searched using k-mer based analysis with overlapping 27-mers identified from the indels and several kb of surrounding sequences from the consensus of the reference genomes. The counts of occurrences of these kmers were smoothed using rolling average in windows of 20 k-mers. The deletions had consistent kmer count of zero throughout their sequences, whereas sequences present in the genomes had positive k-mer counts.

In total we examined 64 species (76 individuals) from Lake Malawi including 7 individuals of *Astatotilapia calliptera*, 88 species (90 individuals) from Lake Victoria and 57 species (69 individuals) from other African lakes or riverine habitats (Supp Table S2). This amounts to 209 species and 235 individuals.

To place these taxa in a phylogenetic context, we use the tree of Meier et al. (2017), which is based on RAD genotypes from the Lake Victoria superflock and outgroup species (Meier et al. 2017, Supp Fig 1). We replaced the Malawi clade in that tree to include the more extensive set of Malawi taxa included here, using a taxonomic tree divided into four Lake Malawi clades: rock, sand, pelagic, and deep (Hofmann et al. 2009) and which includes *A. calliptera* joined with the rock dwelling clade (Joyce et al. 2011, Malinsky et al. 2018). The additional 52 samples genotyped by kmer analysis were grouped by neighbor joining based on distances calculated from a set of SNPs identified across the genome (58 species total; Supp Fig S2; Malinsky et al. 2018). This tree had 12 clades that shared at least one species in common with the RAD tree from Meier et al. (2017). We therefore noted how many of the species examined in this study fall within the clades identified by Meier et al. 2017. In addition, there was one group of samples present in the kmer analyzed genomes which were placed in their own clade because a correspondence could not be made with the Meier et al. 2017 tree.

### Comparison of TE family sizes

Comparisons of TE content requires genomes of high quality built from long sequence reads that can span repetitive sequences. The analysis of specific TE families could only be performed accurately by using long-read based genome assemblies of the Malawi zebra cichlid *Metriaclima zebra* (UMD2a; Conte et al. 2019) and the Nile tilapia *Oreochromis niloticus* (UMD1; Conte et al. 2017). TEs were identified and assigned to families using a combination of RepeatModeler and RepeatMasker. First, RepeatModeler *version open-1.0.8* (Smit and Hubley 2010) was used to identify and classify *de novo* repeat families separately for each assembly. These *de novo* repeats were then combined with the RepBase-derived RepeatMasker libraries (Bao et al. 2015). RepeatMasker *version open-4.0.5* (Smit et al. 2010) was run on the final anchored assembly using NCBI BLAST+ (*version 2.3.0+*) as the engine (‘*-e ncbi*’) and specifying the combined repeat library (‘*-lib*’). The more sensitive slow search mode (‘*-s*’) was used.

### Age of TE insertions

To ground truth the age of one locus, we estimate when the *Rx1* insertion / deletion arose, using sequence divergence for approximately 1450 bp of flanking sequence (i.e. not including sequence for either the deleted or inserted regions). Sequences for 18 Lake Malawi cichlids, *P. nyererei*, *A. burtoni*, and *O. niloticus* were aligned using MAFFT (Katoh et al. 2002, Katoh and Standley 2013). This alignment was analyzed with jModelTest 2.4.1 to determine the best tree model and the optimal analysis parameters (Posada 2009, Darriba et al. 2012). Trees were rooted using *O. niloticus* as the outgroup. The optimal tree and the tree averaged over all the top models were identical in topology. The optimal tree was used to calculate the average distance between the Malawi long palette (high *SWS2A* expression) alleles and either the short or medium palettes (no *SWS2A* expression). The average distance between the Malawi alleles and the *P. nyererei* allele from Lake Victoria was also calculated. To estimate the time when the short and medium alleles diverged from the long allele, we divided the average long to (short, medium) distance by the average Malawi – *P. nyererei* distance and multiplied by the divergence time of species in Lakes Malawi and Victoria, thought to be 2.3 MY (Friedman et al. 2013).

## RESULTS

### Indel analysis

The indel locations in the *M zebra* UMD2a genome are listed in Supp Table S3, with the corresponding sequences given in Supp Table S4. The variation at these loci is shown in Figure 1, where the genotypes of three outgroup species (*A. burtoni, P. nyererei* and *O. niloticus*) are compared to several Malawi cichlid species with known opsin expression. In some cases, a given locus includes either a deletion or an insertion for some individuals.

**Figure 1.**
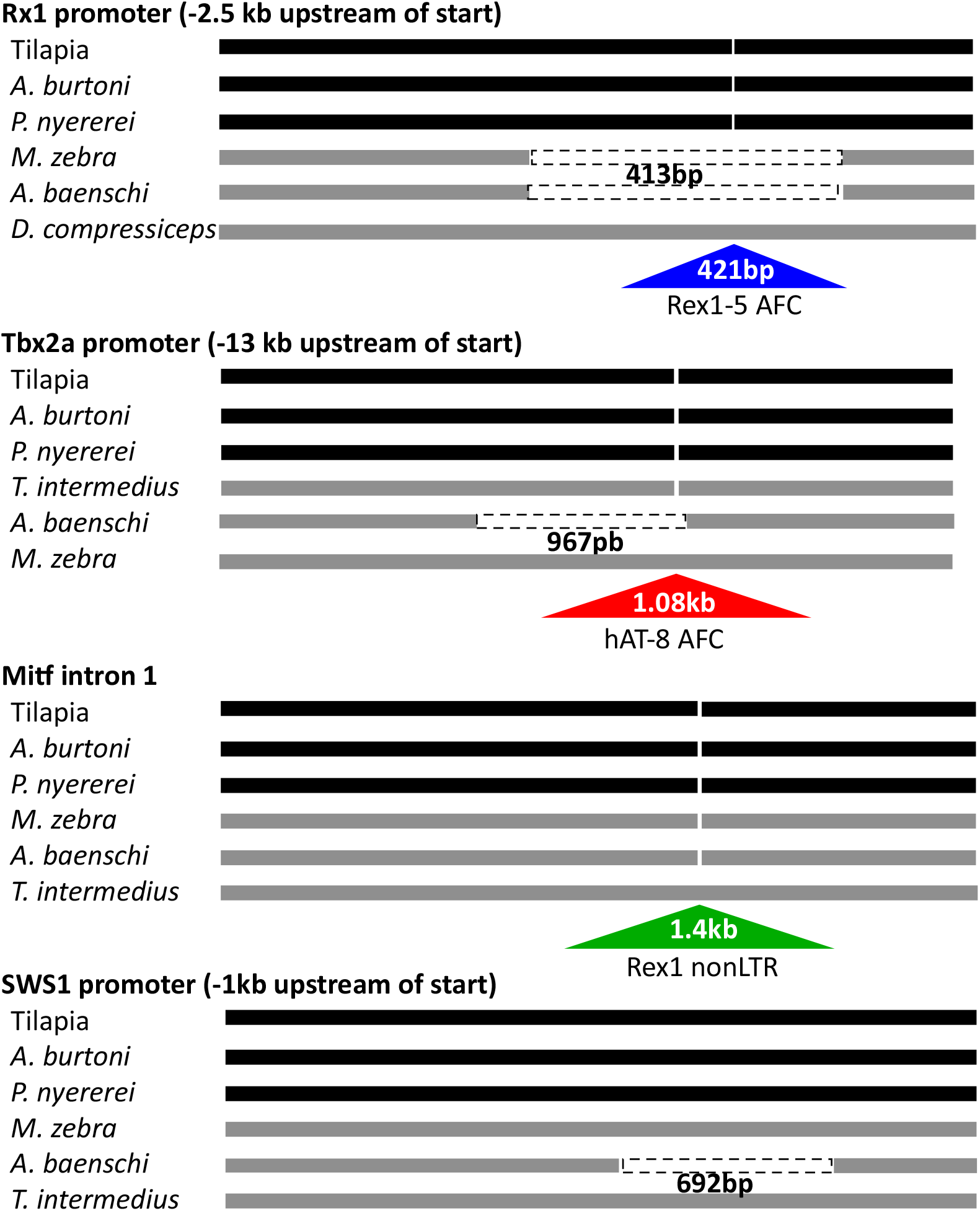
Genomic variation in putative causative mutations underlying changes in opsin expression. Genomic regions for three long palette outgroup species (the tilapia, *Oreochromis niloticus, A. burtoni*, and *P. nyereri* are compared to species from Lake Malawi (*M. zebra* has short palette, *A. baenschi* has medium palette while *D. compressiceps* and *T. intermedius* have the long palette). Deletions are shown as dashed boxes while insertions are shown as triangles.

The *Rx1* locus has either a fixed length deletion (413 bp relative to *P nyererei*) or an insertion of varying length (268-421 bp relative to *P nyererei*). The deletion occurs in short and medium palette LM species and occurs at exactly the same location in each species. The insertion occurs in long palette LM species, but varies in length across species because of its repetitive long microsatellites. Interestingly, the boundaries of the deletion are outside the boundaries of the insertion. We hypothesize that when the deletion occurred, it removed both the insertion as well as some surrounding ancestral sequence. The *Tbx2a* locus includes a 1081bp insertion in *M. zebra* and a 967 bp deletion in *A. baenschi*, relative to *P nyererei*. None of the other species show variation at this location. The *Mitfa* locus involves a 1408bp insertion in intron 1 of the gene and occurs in numerous species. No species with deletions were identified. The *SWS1* promoter involves a 692 bp deletion and occurs only in *A. baenschi*, with no instances of insertions.

The inserted sequences were characterized using Repbase on the Censor website (https://girinst.org/censor/index.php). The longest Rx1 promoter insertions come from *Trematocranus placodon* and *Dimidiochromis compressiceps*. For their 421bp insertions, 384 bp matches Rex1-5 AFC, a known nonLTR retrotransposon. For the 1081bp insertion for the *Tbx2a* regulatory region in *M. zebra*, all but the first 9bp matches hAT-8 AFC, a DNA transposon from the hAT family. The AFC in the names of these repetitive elements indicates they were first described in African cichlids. The last insertion, in intron 1 of *Mitfa*, is a bit more complex. It includes matches to four different transposable elements. The longest match is 522bp of a Rex1 TE from *Petromyozon marinus*. There are several other fragments, which are 50-70 bp long, matching L1 LINES, DNA/Mariner and Copia LTR elements from diverse species. For the final *SWS1* locus, we have not observed any insertions, only the deletion. However, based on annotation of the other three loci, it is possible that a transposable element may have inserted in the SWS1 regulatory region to cause the subsequent deletion.

### Indel age

In order to estimate when the indel might have arisen for one locus, we compared the sequences around the Rx1 indel to date when the two alleles diverged. We built a tree based on the sequence that surrounds the indel region (Supp Fig. 3). Using sequences from Lake Malawi cichlids, *P. nyererei*, *A. burtoni* and *O. niloticus*, it appears that the divergence between the allele with the insertion (long palette) and the alleles with the 413 bp deletion (short and medium palettes) is approximately 25% of the divergence between species in Lake Malawi and *P. nyererei* from Lake Victoria. The Malawi to Victoria split is thought to be approximately 2.3±0.7MY (Friedman et al. 2013). This suggests the allele arose approximately 0.6 MY ago, which is less than the age of LM. While this approximation cannot conclude whether the allele arose within or outside of the Malawi basin, it is comparable in age with early divergences in the Malawi flock.

### Phylogenetic origin

To determine how prevalent these transposable elements are, we searched 64 LM species as well as 145 taxa outside of LM. Using PCR screening we were able to genotype the majority of these loci within LM. Whole genome sequencing provided most of the data for taxa outside of the lake. This included 88 taxa from Lake Victoria proper, 18 species outside Lake Victoria but from the Lake Victoria region superflock (LVRS) and 39 outgroups species. Most of the outgroups were riverine but they included a handful of species from Lake Tanganyika. The complete dataset for the 209 species is given in Table S2 and the data is mapped onto a phylogenetic tree in Figure 2. The major finding from these data is that these indels only occur in LM species, with one exception discussed below. None of the indels are found in any of the species from Lake Victoria or the surrounding region. Three of the indels are not shared by any riverine species. This suggests that these indels are specific to the Malawi flock and have arisen relatively recently, within the ~1MY history of the radiation.

**Figure 2.**
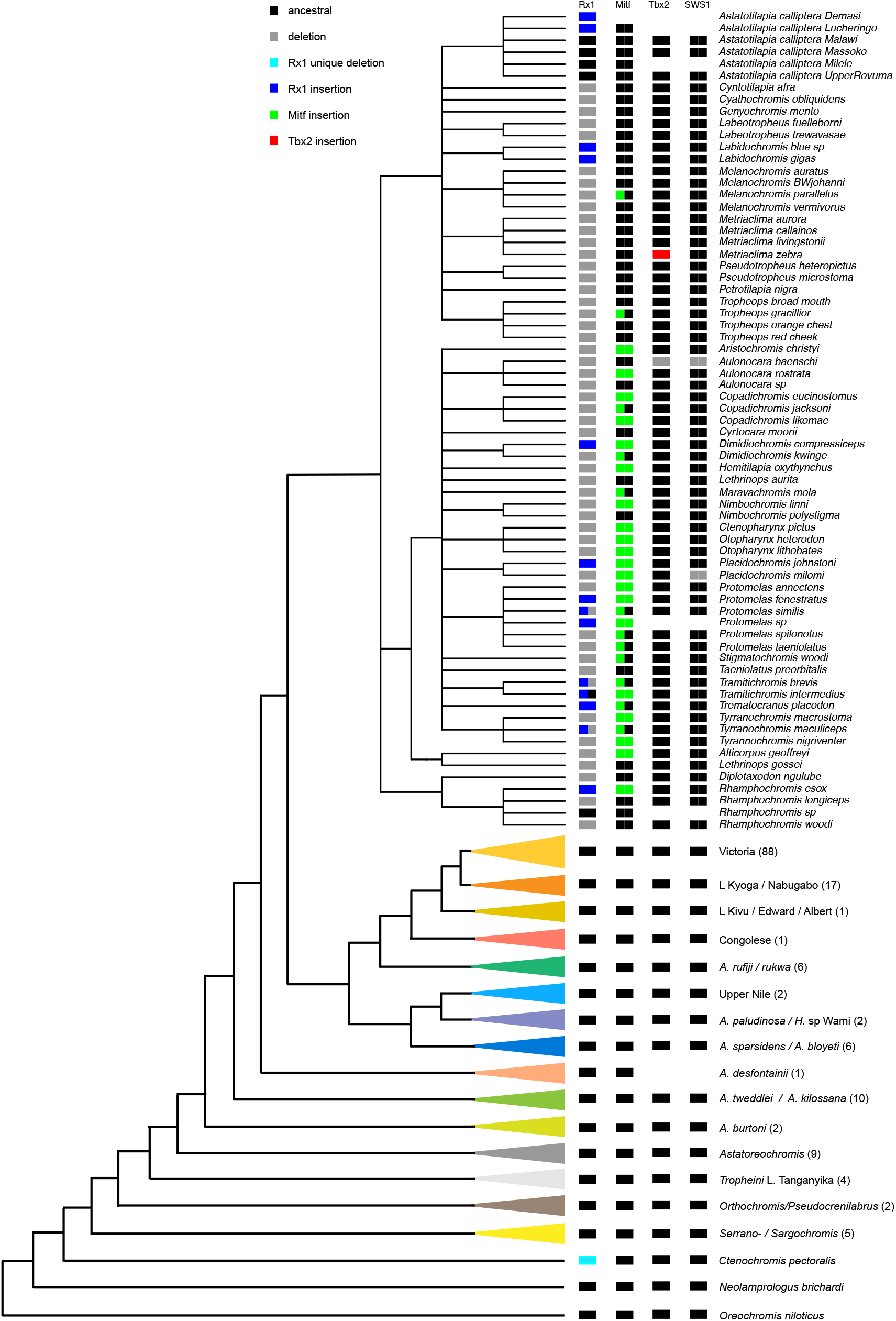
The state of the four indels across the phylogeny of 209 species. This tree is based on Meier et al (2017; Supp Fig 1) with clades color coded as in that original presentation. For species outside of LM, the number of individual sampled in particular clades are noted in parentheses.

The one exception is the indel for the *Rx1* locus. Along with native LM species, the *Rx1* insertion also occurred in *Astatotilapia calliptera* collected from the Lucheringo River. However, *A. calliptera* from rivers nearby Malawi are thought to be closely related and actually phylogenetically imbedded in the Lake Malawi rock dwelling clade, suggesting there is migration in and out of the lake (Malinsky et al. 2018). The *Rx1* deletion was also found in *Rhamphochromis longiceps*. While this species is found in Lake Malawi, our sample came from Lake Chilangali which is only 10km from Lake Malawi. This lineage also phylogenetically groups within the Lake Malawi flock (Figure 2).

There is one additional taxon that has a similar, though distinct, deletion for the *Rx1* locus. *Ctenochromis pectoralis* was sampled from Chemka Hot Springs in northern Tanzania, close to the Kenyan border. This individual has a deletion that overlaps with the LM *Rx1* deletion. On aligning the *C. pectoralis* and *A. baenschi* reads to the *Pundamilia* genome, the *C. pectoralis* deletion length is only 323 bp, instead of the 413 bp found in *A. baenschi* and the other Malawi species. Both edges of the deletion are different, with *C. pectoralis* genome having 94 bp of sequence on one side and 4 bp on the other side that are missing in *A. baenschi*. Because of the physical (~750km) and phylogenetic distance of *C. pectoralis* from the Lake Malawi flock, and the differences in indel boundaries, this must be a unique deletion. This suggests that the *Rx1* regulatory region may be predisposed to deletions.

There is variation in the prevalence of the different indels within Lake Malawi. Of the 76 Lake Malawi individuals examined at the *Rx1* locus, 12 taxa had insertions, 55 had deletions, and 5 had the ancestral sequence matching *P. pundamilia* and *A. burtoni* (this includes multiple *A. calliptera* individuals). In addition, four of the individuals are heterozygous, with three having both an insertion and a deletion allele and one having an insertion plus an ancestral allele. This is in keeping with that fact that the insertion and deletion are quite common across species. The *Mitfa* insertion is also quite common, with 20 individuals being homozygous for the insertion, 12 being heterozygous and 43 having the ancestral sequence (75 individuals total). Again, since we are examining single individuals for most of these species, this means that individuals have one copy of both the insertion and the ancestral sequence. This suggests the insertion is highly prevalent in different species.

In contrast to those two loci, most individuals have the ancestral sequence at the *Tbx2a* and *SWS1* loci (68 individuals total). For the *Tbx2a* locus, only *M. zebra* has the insertion and only *A. baenschi* has the deletion. We examined 4 *M. zebra* individuals and two from Mazinzi Reef had the insertion and two from elsewhere in the lake did not. For the *SWS1* locus, we have not found evidence of any TE insertions and among the taxa included here only two species (*A. baenschi* and *P. milomo*) have the deletion. However, our previous studies which included more taxa found two additional species with the SWS1 deletion (A. stuartgranti and Trematocranus placodon; Nandamuri et al. 2018)).

The TE families that we have identified here include the hAT DNA transposons and the LINE/Rex1/Babar nonLTR transposons. Recent analyses of the highly contiguous PacBio genomes of *M. zebra* and *O. niloticus* indicate that these transposon families are more prevalent in the *M. zebra* genome than in *O. niloticus*. To more broadly examine the prevalence and location of these TEs, we analyze the top three TE families including the Tc1 Mariner and hAT DNA transposons. Genomic locations were divided between 15kb promoters, exons, introns and intergenic regions (Fig. 3; see Conte et al. 2019, Table S5). Very few of the TE insertions occur in exons. For the promoter, intron and intergenic regions, all three TE families have more insertions in *M. zebra* than in tilapia. Obviously, this is linked to the fact that *M. zebra* just has more insertions in total. The one exception is hAT transposon promoter insertions, which occur more frequently in tilapia than *M. zebra*. Even for this class, there are still over 2000 instances of promoter insertions in *M. zebra* suggesting they could have a big impact on gene expression.

**Figure 3.**
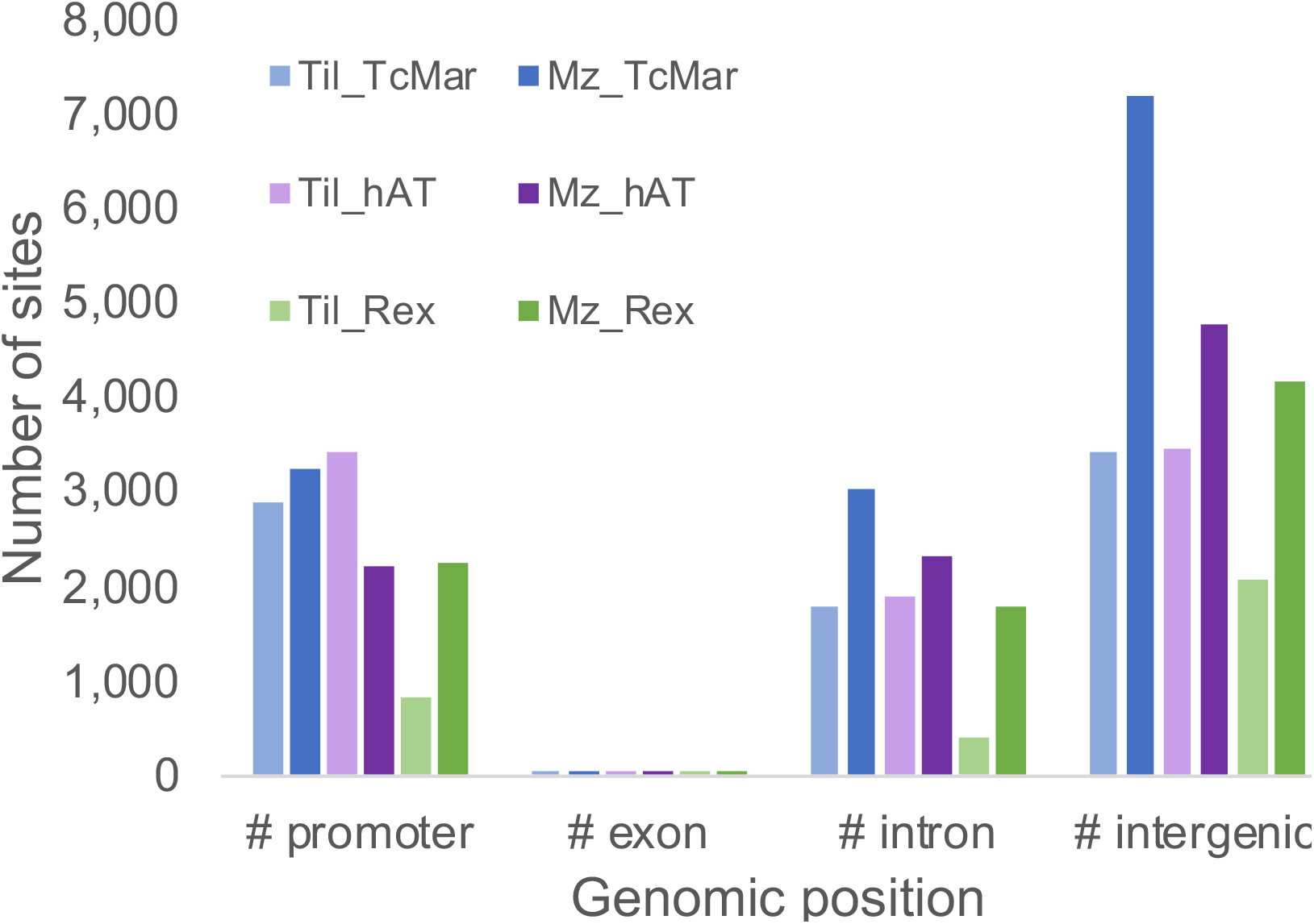
Number of transposable elements from several different families in Tilapia and *M zebra* based on their location. Data provided in Supp Table S5.

## DISCUSSION

This study explored the genetic mechanisms causing differences in a key cichlid phenotype, visual sensitivity. Although the cichlid genome project suggested several possible sources of regulatory mutation, most of these do not seem to play a role in visual system evolution. Coding sequence differences among opsins do not lead to large shifts in retinal cone cell peak sensitivities, which are instead caused by regulatory changes. There is no evidence for cichlid-specific duplications of opsin genes. The regulatory changes do not result from evolution of miRNA target sites in opsin 3’UTRs. Instead, we find three eQTL that correspond to mutations in the promoters of transcription factors that act in trans, along with one eQTL that corresponds to a mutation in the SWS1 opsin promoter. We suggest the movement of TEs can explain this regulatory diversity.

These indels alter gene expression of either the *SWS1* or one of the three transcription factors that alter opsin gene expression in trans. For the *SWS1* opsin gene, the deletion removes a conserved noncoding element (CNE) and a miRNA (Nandamuri et al. 2018). These two regulatory elements are conserved across 230 MY of fish evolution (zebrafish to cichlids). Work in medaka has shown that *SWS1* opsin and the miRNA (miR-729) are expressed in the same photoreceptor and coregulated by the CNE (Daido et al. 2014). Therefore, deleting the CNE would impact *SWS1* and miR-729 expression, with both within the same gene expression network. For Rx1, there is a 413bp deletion that is 2.5kb upstream of the Rx1 start site. This deletion removes several potential transcription factor binding sites (TFBS) including sites for *Tbx2a* and *Mitf*. For *Tbx2a*, there is a 967bp deletion that is 13.4kb upstream of its start site. This deletion removes a probable regulatory region conserved in sticklebacks and medaka (130 MY divergence; timetree.org). It also contains a TFBS for *Rx1*. Finally, *Mitfa* has a 1.4kb insertion in its first intron, which contains a potential *Rx1* TFBS. In this way, these different regulatory regions may work together to switch on various opsins. Therefore, the deletions and insertions will alter the presence / absence of key transcription factor binding sites (Schulte et al. 2014, Sandkam et al. in press).

Our hypothesis for the ultimate molecular mechanism generating this regulatory diversity is that TE insertions either directly alter gene expression (*Mitfa*) or make regulatory regions vulnerable to subsequent deletions (*Rx1, Tbx2a, SWS1*). For two of the deletions, we identified lineages that had TE insertions in the exact same location. We suggest that the excision of the TEs at these sites removed significant regulatory sequence to generate the large (*Rx1:* 413 and *Tbx2a:* 967bp) deletions that we observe in the affected species. Although we did not find a TE insertion associated with the *SWS1* locus, we hypothesize that the large 692 bp deletion may be the result of the insertion and subsequent excision of a mobile TE. The observation that at least three out of four causative mutations underlying eQTLs for opsin expression appear to be associated with TE suggests that TEs are an important factor in changing phenotypes, in this case shifting expression of both transcription factors and their downstream targets.

In trying to date the insertion of these TEs, we find they are recent insertions, present only within species from Lake Malawi. These precise indels were not observed in species outside of the Malawi flock. This includes examination of the sister lineage within Lake Victoria as well as numerous taxa found in the rivers surrounding Lake Malawi. This then bounds their origin to less than 1MY, our best estimate for the age of the flock (Ivory et al. 2016, Malinsky et al. 2018)

One of the loci may show convergent evolution. Examination of the enhancer of *Rx1* suggests that a similar though distinct deletion arose separately outside of the Malawi flock in a phylogenetically distant lineage. This suggests that this regulatory region may be susceptible to TE movement, enabling convergent regulatory mutation.

The idea that TEs might be important in cichlid phenotypes is also supported by an independent previous study. In cichlid pigmentation patterns, egg spots are a unique element that occurs on the anal fin of male haplochromine cichlids. In a study of the genetic basis of cichlid egg spots, Santos et al (2014) identified the gene four and a half LIM domain protein 2 (fhl2b) as important for egg spot formation. The haplochromines with egg spots had a SINE element in the fhl2b promoter that was missing in non-haplochromines, which lacked egg spots. This supports the idea that mobilization of a SINE introduces some new regulatory sequence that introduces the egg spot phenotype.

Previous genome-wide analyses also support the idea of a link between repetitive elements and structural variation in cichlid fishes. The original cichlid genome project found the composition of cichlid genomes to be 16-19% TEs (Brawand et al. 2014) and with long read sequencing the estimates are now 35-37% (Conte et al. 2017, Conte et al. 2019). TE insertions near the 5’UTR were associated with increased gene expression in all tissue types. Another study examined structural variants in these 5 genomes and identified deletions associated with SINE elements and inversions associated with both SINE elements and DNA transposons (Penso-Dolfin et al. 2018). Some of the structural variation was lineage specific.

Transposable elements have been noted to play a key role in evolution (Oliver and Greene 2009). TEs have been proposed to contribute to reproductive isolation and introgression as well as speciation (Serrato-Capuchina and Matute 2018) in a number of groups including amniotes (Zeng et al. 2018), mammals (Ricci et al. 2018), birds (Suh et al. 2018) and fishes (Volff 2005). TE movement is sometimes ascribed to stress in fishes (Symonova et al. 2013, Auvinet et al. 2018). Studies in very young hybrids show that TE number can increase as a result of tandem duplication though they did not find evidence of new insertions (Dennenmoser et al. 2019).

One additional consideration is that TE mobility is thought to be enhanced in hybrids. Since TE movement is repressed by PIWI-interacting RNAs (piRNA), hybridization could lead to an incompatibility of parental piRNAs and the corresponding TEs that allows TEs to increase their mobility (Dion-Cote et al. 2014, Dion-Cote and Barbash 2017, Luo and Lu 2017). Introgression and hybridization have been demonstrated for cichlids in several African lakes (Salzburger et al. 2002, Smith et al. 2003, Meier et al. 2017, Malinsky et al. 2018, Svardal et al. 2019). Therefore, past hybridization events within Lake Malawi cichlids might contribute to the increase in TEs within this flock, some of which could contribute to the genetic and phenotypic diversity we find here. Induced TE movement might be one mechanism for how hybridization contributes to cichlid speciation (Seehausen 2004). Further work would be needed to compare PIWI interacting RNAs between different cichlid species to see whether they have evolved sufficiently to cause mismatches with their target sites.

An important question concerns the relative contributions of positive selection and of genetic drift in the accumulation of the above-described mutations and of other indels linked to TE movement. While estimates of long term effective population sizes (N_e_) in Lake Malawi cichlids are relatively high at 50,000 to 130,000 (Malinsky et al. 2018) or >120,000 (Won et al. 2005), many rocky shore species live in highly structured small populations where current N_e_ may be much smaller and drift much more important. For example, estimated current N_e_ values for species of the rocky shore genus *Tropheops* are 2,000 to 40,000 (Won et al. 2005), while estimated current N_e_ for two rock dwelling species of the genus *Metriaclima* can be as low as 500–1,500 (Husemann et al. 2015). Larger population sampling and population genetic approaches will be required to estimate selection parameters for these indels.

### Conclusions

We have shown that four candidate loci underlying opsin expression are likely the result of insertions and deletions generated by mobile transposable elements. In some cases, we find species with both TE insertions and deletions at the same locus, suggesting that TE movement has caused regulatory indels. These indels are recent and may be an important contributor to cichlid diversity. Indels may result from increased TE movement facilitated by hybridization, with possible fixation in smaller populations due to drift.

## Author contributions

This study was conceived by KC, OS, MM, and TK. Genomic analyses were performed by MC, MM, JM and KC. Laboratory analyses were done by SPN, BS, and SM. All authors contributed to and approved the manuscript.

## Acknowledgements

We thank Richard Durbin and Hannes Svardal for helpful discussions. This study was supported by funding from the NIH (1R01EY024639 to KC), the Swiss National Science Foundation (to OS and 176039 to Walter Salzburger), the US National Science Foundation (DEB-1830753 to TK), and an EMBO grant (ALTF 456-2016 to MM).

