## Supplementary figures for "Movement of transposable elements contributes to cichlid diversity"

### Slide 1
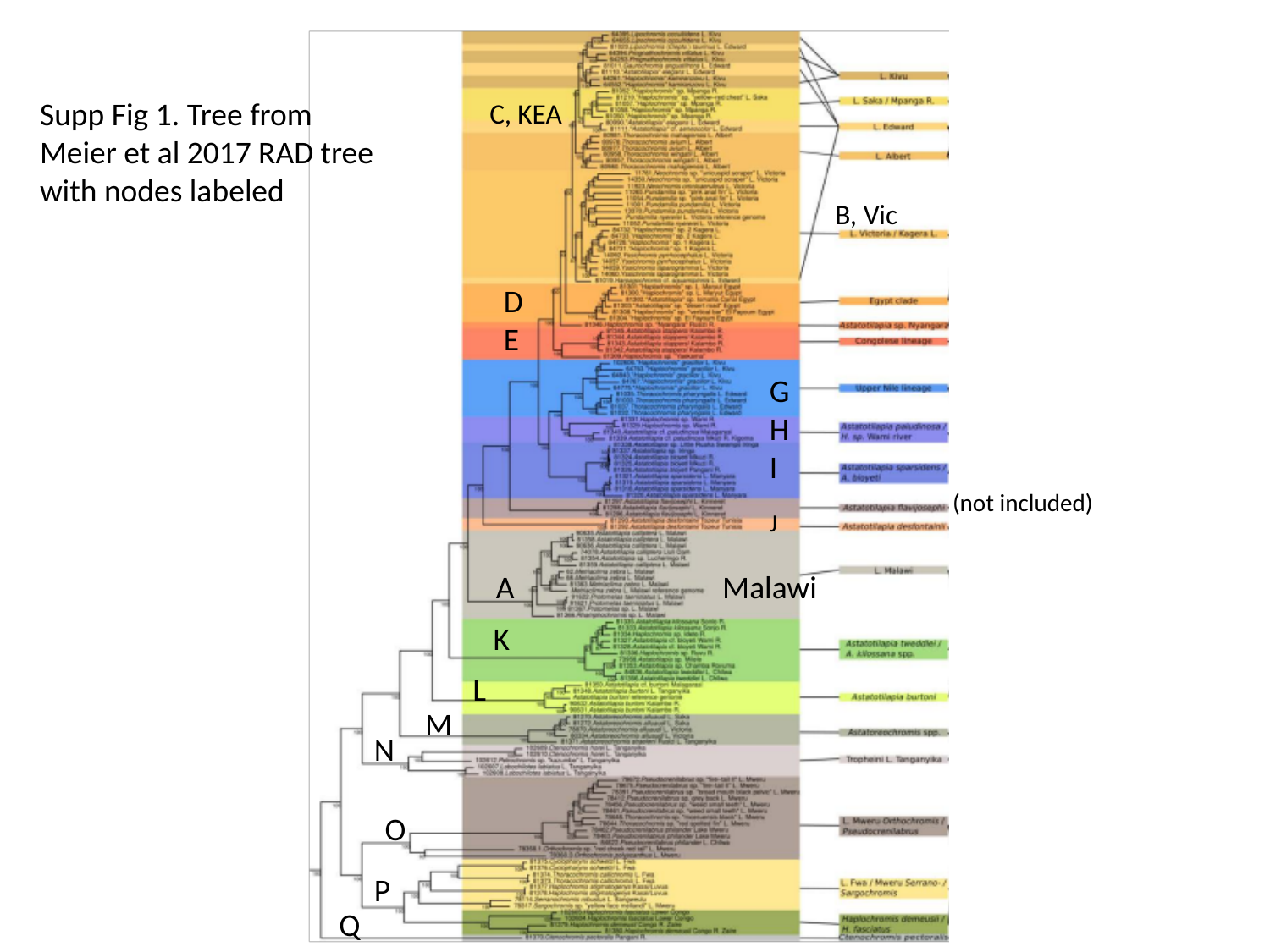

Supp Fig 1. Tree from
Meier et al 2017 RAD tree with nodes labeled
C, KEA
B, Vic
D
E
G
H
I
(not included)
J
A Malawi
K
L
M
N
O
P
Q

### Slide 2
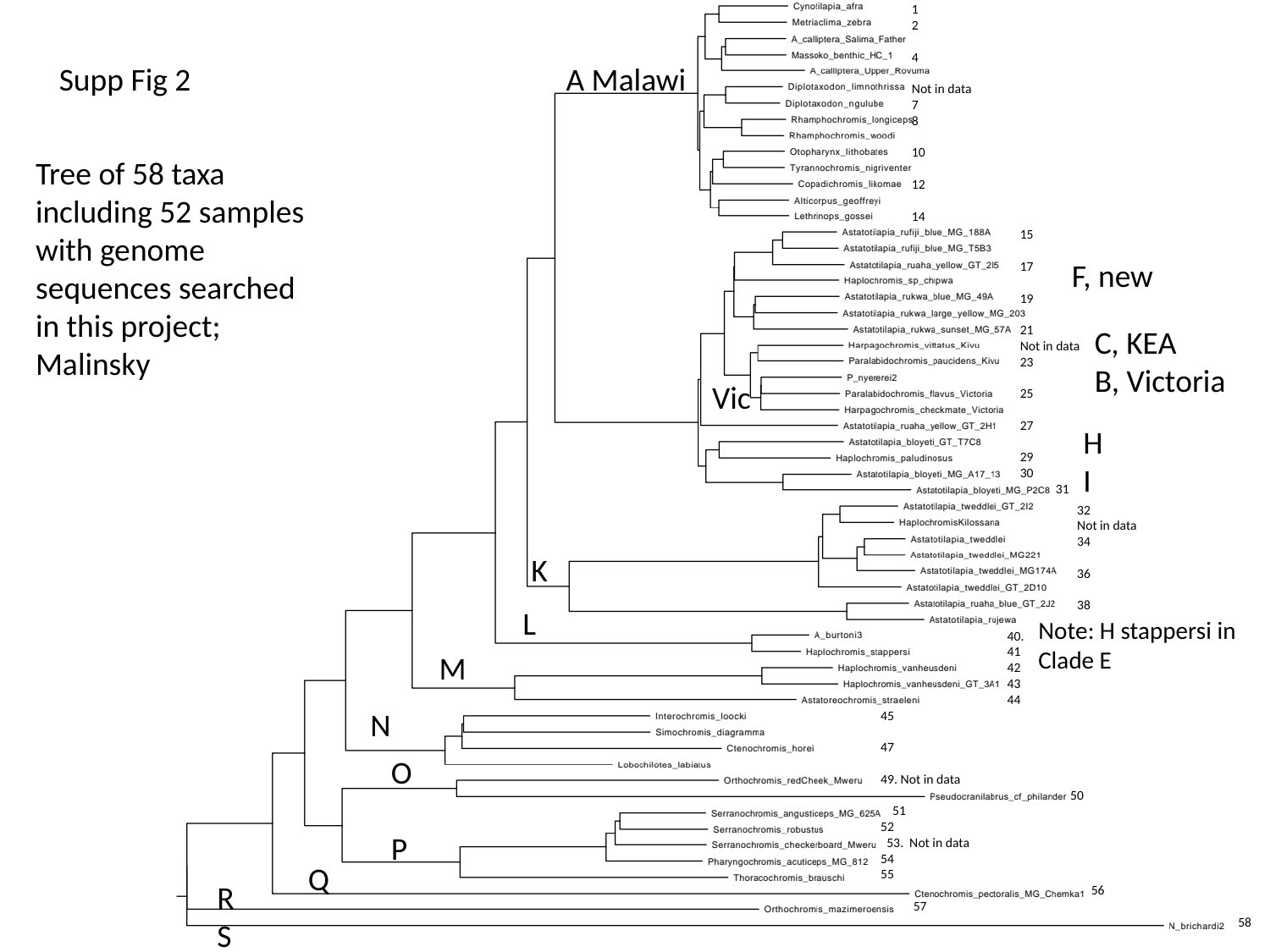

1
2
4
Not in data
7
8
10
12
14
Supp Fig 2
A Malawi
Tree of 58 taxa including 52 samples with genome sequences searched in this project; Malinsky
15
17
19
21
Not in data
23
25
27
29
30
 31
F, new
C, KEA
B, Victoria
Vic
H
I
32
Not in data
34
36
38
K
L
Note: H stappersi in
Clade E
40.
41
42
43
44
M
N
45
47
49. Not in data
 50
 51
52
 53. Not in data
54
55
 56
 57
O
P
Q
R
S
58

### Slide 3
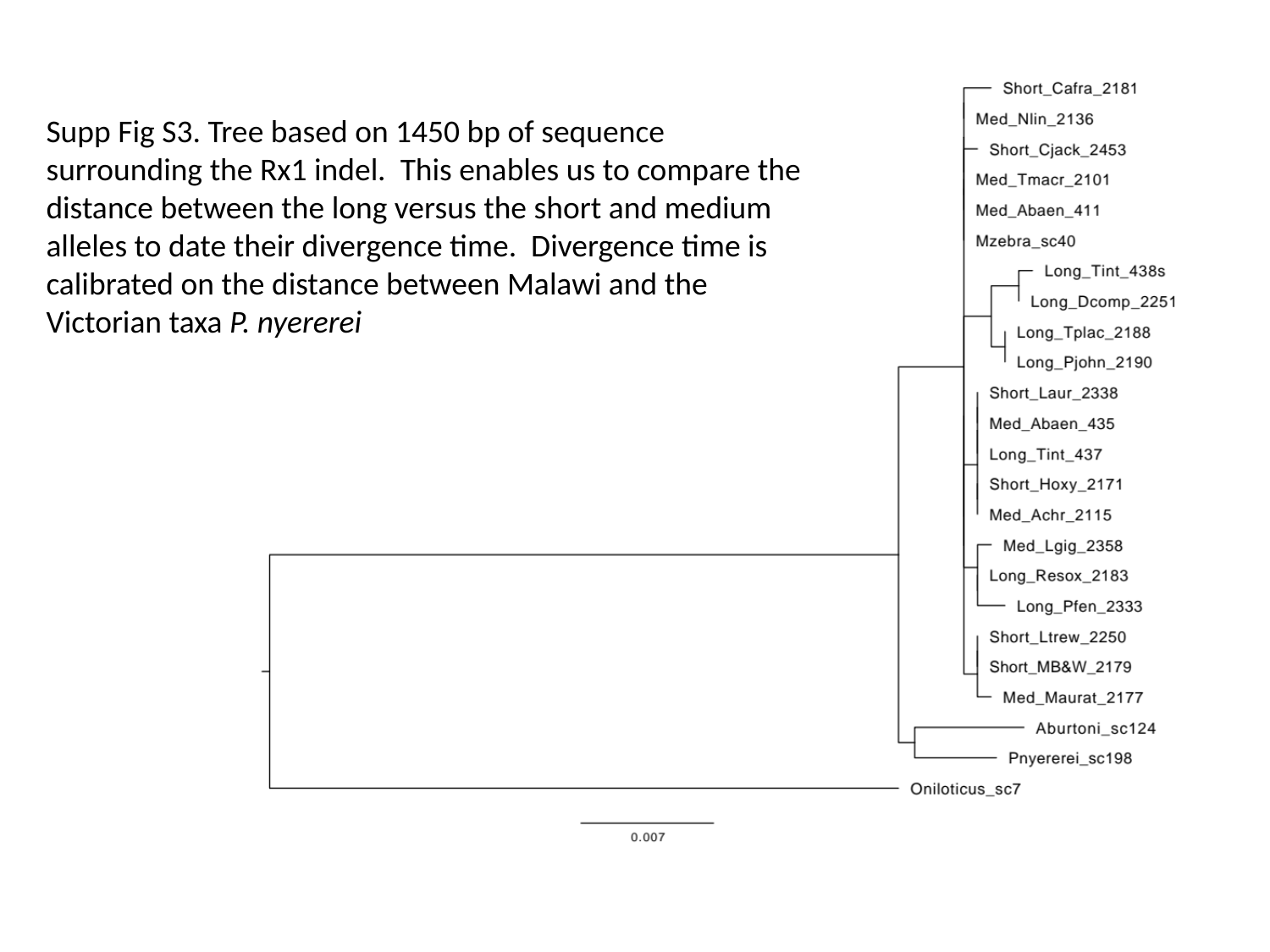

Supp Fig S3. Tree based on 1450 bp of sequence surrounding the Rx1 indel. This enables us to compare the distance between the long versus the short and medium alleles to date their divergence time. Divergence time is calibrated on the distance between Malawi and the Victorian taxa P. nyererei
