## Supplementary tables 1,3,4&5 for "Movement of transposable elements contributes to cichlid diversity"

Supp Table S1. Primers for PCR screening of indels

| Gene |  | Forward |  | Reverse |
| --- | --- | --- | --- | --- |
| Rx1^1^ | Rx1P3F  Rx1KpnF1 | TGGGTTTAATGGCTCCACAG  GCGCGGGTACCCTGCACTCTTTGGCCTTGTT | Rx1P3R  Rx1XhoR1 | CGTTTTCTTTCGCTGGTTACTT  GCGCGCTCGAGTGAGAAGAAACTGTCCCGTGA |
| Tbx2a^2^ | Tbx2_FD-F1 | GGTCAATGATGCTTCCTCTCAG | Tb13kb_R | GAGGTTTGATCAGAGTAAGAACCT |
| Mitf^3^ | MitfP6F  MitfP7F | TGTGTTTTGGGTGTTCAGAACA  GTTGGACAGGTGCTGAAATGT | MitfP6R  MitfP7R | ACGTGCTTGCTTTGCAACAT  ACATCGTCGTTGTAGCTTGAC |
| SWS1^4^ | mnpromoterF | CCTCAATGCTCCAAAAAGGA | promoter_3R | GCCAACCAGCTCACTGTTGT |

^1^Schulte et al 2014

^2^Sandkam et al 2019

^3^Nandamuri 2018

^4^Nandamuri et al 2018

Table S3. Location of regulatory sequences in *Metriaclima zebra* UMD2a genome (Conte et al )

| Nearby gene | Cross | Region | Indel location in *M zebra** | Start site of gene and  NCBI Accession # | M zebra status | *A baenschi* status | Other cross parent status |
| --- | --- | --- | --- | --- | --- | --- | --- |
| Rx1 | TA | Regulatory region 2.5kb upstream | LG23: 1374765<> 1374766 | LG23: 1377294  XM_004555105.1 | Deletion | Deletion | Insertion heterozygous,  *T. intermedius* |
| Tbx2a | TA | Regulatory region 13kb upstream | LG10: 8403573..8402302 | LG10: 8389309  XM_004557464.1 | TE insertion | Deletion | Ancestral |
| Mitfa | TA | Intron 1 | LG5: 7206112<>7206113 | LG5: 7225844  XM_024802633.1 | Ancestral, no insertion | Ancestral | Insertion, *T intermedius* |
| SWS1 | MA | Regulatory region upstream of SWS1 | 000578F_pilon_quiver: 239435..240124 | 000578F_pilon_quiver: 240877  NM_001310074.1 | Ancestral, no deletion | Deletion | Ancestral, *M. ‘mbenjii* |

* Regions deleted in other species listed from start to end. Location of insertion is marked with <>

Supp Table S4. Sequences for four genomic indel regions taken from *M. zebra* genome UMD2a (Conte et al 201x). For the SWS1 and Tbx2a loci, this lists the sequence that is present in *M. zebra* but missing in *A. baenschi*. For the Mitf locus, it denotes the location of the insertion in different species including *T. intermedius*. For the Rx1 locus, it shows the location of the deletion which is shared by *M. zebra* and other short or medium palette species.

>MzUMD2a_SWS1pro_deleted_in_Abaenschi

ACAGAAACCTAACATTAAACAGTGAAAGCAGCAGACATAATCTCCGGTCTTCATACCTGAACCACCAGGAGCTTAATCAATAATCCATAGTAGCAAGTAATCAATAGTCAATACATCAGTGATTAATCAGTCTTTTAAGGCTGCAAGTCACCTCCAAACTGAAGTCGTATCATACCCATGCTCCATGCAGCTCACAGCCTGGTTATGATACAGCCCCCGCTCTAAAGGTCAGCAATCATTACAGCCATCGATCAATCGATCAGTCAATAAATGTGTCTCCTCTTCAGTCTGGTCCAGGATGAAAATGGGCAACAGAAGTGGAGACAGTGCAGTTTCTTCTTGTCCGCGACGTTCTCCGGACTCTGAGCAGGAGAAGTGATGAAGGTTGGAGGACAGCTCCTTCTTAAAGGCTCAGTGTTCTTCCTCCTCCTCTTCCTCACAGTCTTCCTCACGCTGAACCCCTTTAACCGTTTCAGTGGTCGTGAGTCTTCTAATCTGACTGTGTGACGATGTTTTAAGGATTTGGAGGATTGAGGAGGATCACCTGGTCAGGTAAATCTGAAATATCCGGATTACATCGGAAGTTGAGCACACGGAAAAACAAAAGACTCTTATTGGATTTAGATCCGTCAGCCACCTGCTGCTGCTCTTCATCATCAGGCGTCTTCATCGCCCTGCAGTGGGCCTGAC

>MzUMD2a_Tbx2aEnhancer_deleted_in_Abaenschi

AAAAAACTATTTATGCATTAATTACTTCATTTCACAGTGTCTAGACTGTTTGTTGAACTTGCCGCAACAGACAATTTGAAATTACAGCCAGCTTGGGAGTAAGTGATGTGACCCAAGATTGGAGCAGGATTTAGAGGTATGGTGAGTAGGGCTGGGTATCGTCACTGATTTCTAGAATCGATTCAATTCCGATTCACAAGGTCCCGAATCGATTCGATCCACGATTCGATTTAATTCGATTCGATTCGATTTAAATCTGGGAAATTTTGACAGTCAGAGATATAATTCAGATCAGGACATTTACATATTTTTGTATCTATAAAAAGGAAGCTGACACTCTCAAGACTTTATCCAAGGTGTGAGCATCACAGCAGATGCCTTTGTGTCAAAGTAGCTGAAGATAAAACACAGAAAAACATGTAGGTGATTTTCCTGGCCTGGGATTTTATAAAAATATTCTGCAGTACATCAAAAACGAAAGAAAACCATTAATTAATCAACATTACCTCTGACGTTACAGCGGTTTTATTAGAGACATGGCTAAGCATTTTGCATTTTGAATAATTTTAAAAAGTTTCAATTTGTTCAGTATTGAACAGCAGAAATGAGGTTTTCTTTTCGGAAGAATGTAAAAGGGGAAAAAACAGCGGCCGACAGCGCTGTAAGCAACAGTAGACTTGTGTAACAAGCAAGCGAATAATGAAGAAAGCGAAACTGTTCGAGAGAGAGAGAGAAAGGGAGAGGGAGAGAGAGAGAGCTGCGCATTGCAATGGAAGCAAAACAGTAAAAGAGTGAATTCATGACGATGTTTATGTGAAGCGTTTGGATCTTCTTTTGCTGCTGATTCGGTCGATATTGTTTGGAGAGAGATCAAACTAACAGCTTTAGAATCAGCGCATAAAGGGCGTGAACACAAAGCGCGGACCCGGATCAGCGAGCTGTCGGCTTTCAGCCCCGACTGTGAGAAAGGCGACATCTAACTGATTCTGATCCGCGGGTTGCGCTGTGTGTTTAAGTCTATGTGCAGAATCCCTGTACCTGCTCTCATTTACTGTTTGGTGGTTCTTGAAATTTTGTGAGATGTCACCAGAGATTCTGGCATTTCGGACAAAATAAATTTATATTAAAAAATCGATTCAGGATTTTAATGAATCGATTTTATGTTATCCAAGCCAGAATCGATTTTAATCGATGAATCGATTATAAAAACCCACCCCTAATGGTGAGGTTAAGATTCTTAACTGTCTCTAAGTCCTTTGATTGTTTACCTGTAAAATCTTTTTCACCTCACACACATTACGAACAGTTTGAAGGCAGATTGACCTGCCCAGG

>MzUMD2a_Mitf_intron1_Tramitichromis_insertion

GAGTAGATGCGATTTTTACACTAAAAGAGTCAAAATATCTTTGCTGCGCTTGAAAACAGGACTTAAGGTTGAGGATTAGGGTTGAGTTCTAAATACTTCATTATGTGTACCCAGTTCACAAGTCTTTGGTTAAATGAAGACAGGGAAGCTGTTTTACACACAGTCGTATTTAAATTGCTGCACGAGTTGGACAGGTGCTGAAATGTTATATAGTGCTCTTTTAGAAAAAAAGAGCTTGATGAAAAAAAATGTAAAAGAAATGTTGTTTGGACATTTCAACATCTTATCACCGTA<insertion>ATGTTGCAAAGCAAGCACGTTAGATTATTTGGTTTTTAAAAGGCTGTTTTCAGGGGAACTTTTAATGTAACAGAGTAGTATAGTGTGATTAATTTGGATTTGTAAGTTGCACTTTGAAGCTGGAAACATTTGAACATTTAAGTTCTTGCATTTTTAATTTGTGCTTTTACCACACCGGGACAGATGAATTATTATAATGAAAACATTCATTTTTA

>MzUMDa_Rx1pro_deleted_in_short_medium_palettes

CTGCATCCTCGGGCAAAATGCTGAACCCTACGTTTCCCCTGATGCATTTGAGTGAGTGTGTGAATGTGATTGTATGAGTGGGTGAATGAAAAGATGCTAT<deletion_here>TTTACTGCGGGGCATGTGCAGAAAAAATACAGTCCTTTGTAGCACAGAAGTGTAACATAAAAACAAAAAGACTGTGACGAGGAATTTGAAATGATCACGGGACA

1377294

Table S5. Frequency and location of recent insertions for three TE families.

| TE_family in assembly | # insertions (div < 2.0) | #total length of insertions, div < 2.0) | % promoter | % exon | % intron | % intergenic | # promoter | # exon | # intron | # intergenic |
| --- | --- | --- | --- | --- | --- | --- | --- | --- | --- | --- |
| DNA/TcMar in O_niloticus_UMDNMBU | 8,127 | 7,470,032 | 35.73 | 0.32 | 21.94 | 42.01 | 2904 | 26 | 1783 | 3414 |
| DNA/TcMar in M_zebra_UMD2 | 13,467 | 13,922,869 | 24.07 | 0.09 | 22.51 | 53.32 | 3242 | 12 | 3031 | 7181 |
| Mz - On | 5,340 | 6,452,837 | -11.66 | -0.23 | 0.57 | 11.31 | 338 | -14 | 1248 | 3766 |
| Ratio Mz / On | 1.66 | 1.86 | 0.67 | 0.28 | 1.03 | 1.27 | 1.12 | 0.47 | 1.70 | 2.10 |
| DNA/hAT in O_niloticus_UMDNMBU | 8,785 | 3,839,634 | 38.80 | 0.25 | 21.43 | 39.51 | 3409 | 22 | 1883 | 3471 |
| DNA/hAT in M_zebra_UMD2 | 9,316 | 6,931,987 | 23.78 | 0.12 | 24.79 | 51.32 | 2215 | 11 | 2309 | 4781 |
| Mz - On | 531 | 3,092,353 | -15.02 | -0.13 | 3.36 | 11.81 | -1193 | -11 | 427 | 1310 |
|  | 1.06 | 1.81 | 0.61 | 0.48 | 1.16 | 1.30 | 0.65 | 0.51 | 1.23 | 1.38 |
| LINE/Rex in O_niloticus_UMDNMBU | 3,331 | 1,946,563 | 25.01 | 0.51 | 11.86 | 62.62 | 833 | 17 | 395 | 2086 |
| LINE/Rex in M_zebra_UMD2 | 8,219 | 6,291,320 | 27.44 | 0.21 | 21.55 | 50.81 | 2255 | 17 | 1771 | 4176 |
| Mz - On | 4,888 | 4,344,757 | 2.43 | -0.30 | 9.69 | -11.81 | 1422 | 0 | 1376 | 2090 |
|  | 2.47 | 3.23 | 1.10 | 0.41 | 1.82 | 0.81 | 2.71 | 1.02 | 4.48 | 2.00 |
